# Indoxyl sulfate, a gut microbiome-derived uremic toxin, is associated with psychic anxiety and its functional magnetic resonance imaging-based neurologic signature

**DOI:** 10.1101/2020.12.08.388942

**Authors:** Christopher R Brydges, Oliver Fiehn, Helen S Mayberg, Henry Schreiber, Siamak Mahmoudian Dehkordi, Sudeepa Bhattacharyya, Jungho Cha, Ki Sueng Choi, W Edward Craighead, Ranga R Krishnan, A John Rush, Boadie W Dunlop, Rima Kaddurah-Daouk, for the Mood Disorders Precision Medicine Consortium

**Author notes:** Corresponding Authors Rima Kaddurah-Daouk, PhD, Duke University Medical Center, DUMC 3903, Blue Zone South, Durham, NC, USA, Boadie Dunlop, MD, Emory University College of Medicine, 12 Executive Park Dr. NE, Room 347, Atlanta, GA 30329, USA.

## Abstract

**Background:** It is unknown whether indoles, metabolites of tryptophan that are derived entirely from bacterial metabolism in the gut, are associated with symptoms of depression and anxiety.

**Methods:** Serum samples (baseline, 12 weeks) were drawn from participants (n=196) randomized to treatment with cognitive behavioral therapy (CBT), escitalopram, or duloxetine for major depressive disorder.

**Results:** Baseline indoxyl sulfate abundance was positively correlated with severity of psychic anxiety and total anxiety and with resting state functional connectivity to a network that processes aversive stimuli (which includes the subcallosal cingulate cortex (SCC-FC), bilateral anterior insula, right anterior midcingulate cortex, and the right premotor areas). The relation between indoxyl sulfate and psychic anxiety was mediated only through the metabolite’s effect on the SCC-FC with the premotor area. Baseline indole abundances were unrelated to post-treatment outcome measures, which suggests that CBT and antidepressant medications relieve anxiety via mechanisms unrelated to gut microbiota.

**Conclusions:** A peripheral gut microbiome-derived metabolite was associated with altered neural processing and with psychiatric symptom (anxiety) in humans, which provides further evidence that gut microbiome disruption can contribute to neuropsychiatric disorders that may require different therapeutic approaches.

## INTRODUCTION

The gut microbiota impacts numerous aspects of human health and disease (1), including neuropsychiatric disorders. The “microbiota–gut–brain axis” refers to a bidirectional communication pathway that connects the central nervous system (CNS), the gut, and the microbial community that inhabits the gastrointestinal tract (2). Within this axis, the gut microbiota modulates central processes through the activation of neuronal pathways (e.g., the vagus nerve) as well as through the production of microbial metabolites and immune mediators that can trigger changes in neurotransmission, neuroinflammation, and behavior (3-6).

Disruptions to the gut microbiome have been correlated with several neurological disorders, including Parkinson’s disease, autism spectrum disorder, schizophrenia, and major depressive disorder (MDD) (7-10), though the specific mechanisms that underlie the role of the gut microbiota in these diseases is not fully understood. However, research in preclinical rodent models shows that the gut microbiota is sufficient to alter host behavior, as shown by the increase in anxiety- and depressive-like behaviors in rodents after fecal microbiota transfer from humans with depression relative to those that received transfer of fecal microbiota from demographic controls (11-12). Further, transferred microbes resulted in altered metabolic states in the recipient mice that displayed depressive-like symptoms (12). These data implicate the gut microbiota as direct contributors to behaviors associated with depression and anxiety through their metabolic effects. In this study, we explore gut microbiota-associated tryptophan metabolism and correlate levels of metabolites to clinical symptoms and severity of depression and anxiety in humans.

Tryptophan is an essential amino acid that can be metabolized in the gastrointestinal tract via the serotonin, kynurenine, and indole metabolic pathways (Figure 1), which have been associated with human maladies including autoimmunity, inflammatory diseases, metabolic syndrome, and neurological diseases including depression and anxiety disorders (13,14). Strikingly, the gut microbiota is exclusively responsible for the conversion of tryptophan in the indole pathway, as there are no detectable levels of indole or indole derivatives in gnotobiotic mice that lack a gut microbiome (15). Analysis of biosynthetic pathways found that the genes necessary to make indole and indole derivatives, such as indole-3-propionic acid (IPA), indole-3-acetic acid (IAA), and indole-3-lactic acid (ILA), are found exclusively in the gut microbiome but not in mammalian genomes (13) (Figure 1). These indoles can have important immunomodulatory effects and are potent agonists for aryl hydrocarbon receptors (16) (AHRs), which regulate host immunity and barrier function at mucosal sites (17).

**Figure 1.**
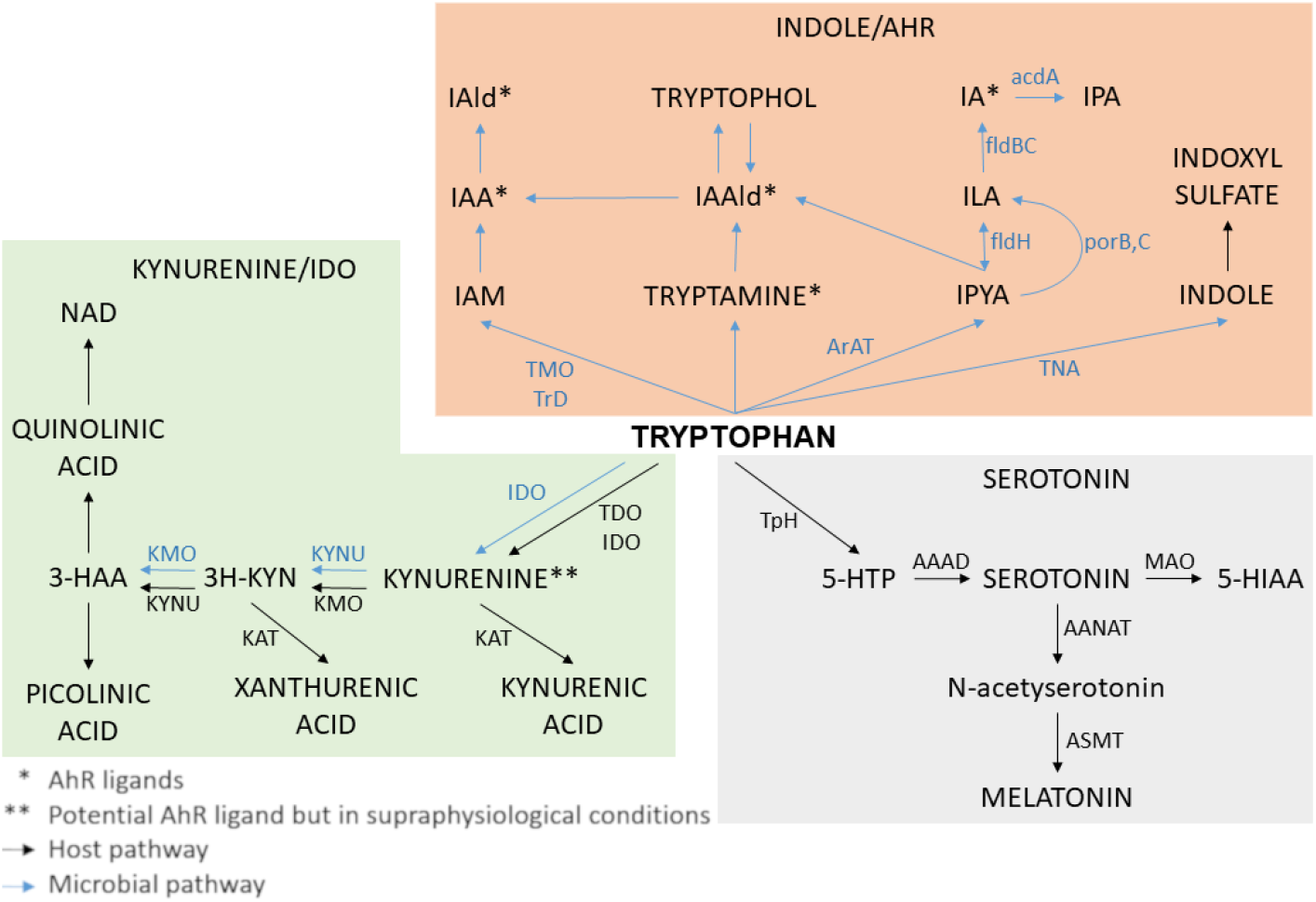
Tryptophan human gut bacterial co-metabolism leading to production of indoles including IPA, IAA, ILA and IS. *Abbreviations*: 3-HAA: 3-Hydroxyanthranilic Acid; 3H-KYN: 3-Hyroxykynurenine; 5-HTP: 5-hydroxytryptophan; AAAD: Aromatic Amino Acid Decarboxylase; AANAT: Aralkylamine N-Acetyltransferase; acdA: acyl-CoA dehydrogenase; AraT: Aromatic Amino Acid Aminotransferase; ASMT: Acetylserotonin O-Methyltransferase; fldBC: phenyllactate dehydratase; fldH: phenyllactate dehydrogenase; IA: Indole Acrylic acid; IAA: Indole Acetic Acid; IAAld, indole-3-Acetaldehyde; IAld: Indole-3-Aldehyde; IAM: indole-3-Acetamide; IDO: Indolamine 2,3-Dioxygenase; ILA: Indole-3-Lactic Acid; IPA: Indole-3-Propionic Acid; IPYA: Indole-3-Pyruvate; KAT: Kynurenine aminotransferase; KMO: Kynurenine 3-Monooxygenase; KYNU: Kynureninase; MAO: Monoamine Oxidase; NAD: Nicotinamide Adenine Dinucleotide; porB, C: pyruvate : ferredoxin oxidoreductase B and C; TDO: Tryptophan 2,3-Dioxygenase; TMO: Tryptophan 2-Monooxygenase; TNA: Tryptophanase; TpH: Tryptophan Hydroxylase; TrD: Tryptophan Decarboxylase.

Indole derivatives can also affect immune status in the brain, as some indole derivatives (e.g., IPA and IAA) have anti-inflammatory effects on neurodegenerative diseases in the experimental autoimmune encephalomyelitis (EAE) mouse model of multiple sclerosis (18,19) as well as in a cell line model of Alzheimer’s disease (20). Other indole derivatives can be further metabolized by host processes into molecules that may be harmful to human health. Specifically, indole can be sulfonated in the liver into uremic toxin indoxyl sulfate (IS), which crosses the blood-brain barrier (21) (Figure 1). IS, which is normally cleared via the kidneys and excreted in the urine, is associated with cardiovascular disease in patients who have chronic kidney disease via induction of oxidative stress in endothelial cells (22), and peripheral IS concentrations are associated with diminished cognitive function in renal dialysis patients (23).

IS is also associated with both neurodevelopmental and neurodegenerative diseases, as levels of IS are increased in patients who have an autism spectrum disorder (24) or Parkinson’s disease (25). Although the mechanistic role of IS in these diseases is unknown, IS increases levels of oxidative stress and pro-inflammatory cytokine signaling in astrocytes and mixed glial cells during *in vitro* administration (26), which suggests that inflammation and reactive oxygen species may be involved. Further, IS has been associated with behavioral defects in preclinical models of anxiety and depression. The administration of IS into rodents’ drinking water results in increased concentrations of IS in the brain and increased blood-brain barrier permeability in an AHR-dependent manner, with accompanying increases in anxiety and cognitive deficits (27,28). Monocolonization experiments with indole-producing *Escherichia coli* and isogenic mutants have shown that indole production by gut bacteria is sufficient to drive increases in anxiety- and depressive-like behavior in rats (29).

Taken together, the preclinical and clinical data indicate that indole derivatives provide excellent models to study the microbiota-gut-brain axis given their connection to central immune regulators (i.e., AHR), their link to human neurological diseases, and the exclusivity of indole production to gut microbes.

To date, the effects of peripheral metabolic concentrations on neural functioning have received little study, likely due to the paucity of datasets that contain concurrently collected metabolomic and neuroimaging measures. Such research is crucial for determining how changes in peripheral systems may yield alterations in brain function that can produce clinically relevant symptoms such as depression, anxiety, or cognitive impairment.

Using blood samples stored from the Prediction of Remission in Depression to Individual and Combined Treatments (PReDICT) study, which was a large study of treatment-naïve patients with MDD, we measured levels of four indole derivatives (IPA, IAA, ILA, IS) to address the following questions:

1. Do levels of indoles and their ratios at baseline prior to treatment correlate with depression and anxiety severity at baseline?
2. Do levels of indoles and their ratios at baseline correlate with specific individual symptoms of depression?
3. Can symptom change after treatment with duloxetine, escitalopram, or cognitive behavioral therapy (CBT) be predicted by baseline levels of indoles, and does symptom change correlate with changes in levels of indoles after treatment?
4. Are there relationships between baseline peripheral metabolic concentrations of indoles and brain resting state functional connectivity as determined using functional Magnetic Resonance Imaging (fMRI).

## MATERIALS AND METHODS

### Study Design

The PReDICT study protocol (30), clinical results (31) and initial neuroimaging analyses (32) have been published previously. The study was conducted through the Mood and Anxiety Disorders Program of Emory University from 2007-2013. The study was approved by Emory’s Institutional Review Board. All patients provided written informed consent to participate.

PReDICT was designed to identify predictors and moderators of outcomes to three randomly assigned first-line treatments for MDD: duloxetine, escitalopram, or CBT. The study enrolled treatment-naïve adult outpatients, aged 18–65 years, who had current MDD without psychotic symptoms. To be eligible for randomization, participants had to score ≥18 at screening and ≥15 at baseline on the HAM-D. Key exclusion criteria included the presence of any medically significant or unstable medication condition that could impact study participation, safety, or data interpretation; any current eating disorder, obsessive-compulsive disorder, or any current substance abuse or dependence. Treatment was provided for 12 weeks with duloxetine (30-60 mg/day), escitalopram (10-20 mg/day), or CBT (16 individual one-hour sessions).

### Symptom Assessments

At the baseline visit, participants were assessed by trained interviewers using the HAM-D and the HAM-A. The HAM-A is a 14-item measure that consists of two subscales, “psychic anxiety” (items 1–6 and 14), and “somatic anxiety” (items 7–13) (33). Psychic anxiety consists of the symptoms of anxious mood, tension, fears, depressed mood, insomnia, impaired concentration, and restlessness. Somatic anxiety consists of physical symptoms associated with the muscular, sensory, cardiovascular, respiratory, gastrointestinal, genitourinary, and autonomic systems. Participants also completed the QIDS-SR, which assesses the nine diagnostic symptom criteria for MDD (34). The HAM-D, HAM-A, and QIDS-SR were repeated at the Week 12 visit.

### Blood Sampling

Participants who met all eligibility criteria at the baseline visit underwent an antecubital phlebotomy, without regard for time of day or fasting/fed status. Sampling was repeated at the week 12 visit. Collected samples were allowed to clot for 20 minutes and then centrifuged at 4°C to separate the serum, which was frozen at -80°C until being thawed for the current analyses.

### Neuroimaging

To explore associations between indole metabolites and brain function, we used the resting state fMRI scans collected during the week prior to baseline, the details of which have previously been published (32). Briefly, eyes-open scanning was performed for 7.4 minutes in a 3-T Siemens TIM Trio (Siemens Medical Systems, Erlangen, Germany). Echo planar images were corrected for motion and slice-time acquisition. Scans with head motion >2 mm in any direction were removed from the analysis. The nuisance regressors, including head motion parameters, signal from the ventricle mask, and signal from a region of local white matter, were cleaned. Subsequently, data were applied a band-pass filter and smoothed using an isotropic Gaussian kernel of 8 mm full width at half maximum. The imaging anatomical and functional data sets were co-registered and normalized to standard Montreal Neurological Institute (MNI) 1-mm voxel space. Image analysis was conducted using Analysis of Functional NeuroImages (AFNI) (35,36) 3dvolreg. Consistent with our prior analyses (32), we used a region-of-interest seed-based approach to assess the resting state functional connectivity (RSFC) of the SCC. The SCC volume was defined using the Harvard-Oxford Atlas (37), and the SCC was thresholded at 50% probability centered on MNI coordinates 66, 24, –11. The seeds comprised two 5-mm radius spheres, with a final volume of 485 mL each. Utilizing 3dNetCorr (38), the mean time course of the bilateral seed was correlated voxel-wise with the rest of the brain. The voxelwise correlation coefficients were then z-scored by calculating the inverse hyperbolic tangent, yielding the seed-based RSFC maps for analysis.

### Metabolomics Data Acquisition

Metabolomics data focused on primary and polar metabolites using gas chromatography – time of flight mass spectrometry (39). Briefly, 30 μl of plasma was extracted at -20°C with 1 mL degassed isopropanol/acetonitrile/water (3/3/2). Extracts were dried down, cleaned from triacylglycerides using acetonitrile/water (1/1), and derivatized with methoxyamine and trimethylsilylation. Samples (0.5 μL) were injected at 250°C to a 30 m rtx5-SilMS column, ramped from 50-300°C at 15°C/min, and analyzed by -70 eV electron ionization at 17 spectra/s. Raw data were deconvoluted and processed using ChromaTOF vs. 4.1 and uploaded to the UC Davis BinBase database (40) for data curation and compound identification (41). Result data were normalized by SERRF software to correct for drift or batch effects (42).

### Statistical Analyses

Indole abundance and ratios of each indole pair were included in all analyses. In order to investigate the role of indoles in depression and anxiety symptomology at baseline, partial Spearman rank correlations were conducted between the baseline abundance/ratio of each indole and HAM-D 17-item total score, HAM-A total score, HAM-A Psychic and Somatic subscores, QIDS-SR 16-item total score, and each individual QIDS-SR item after accounting for age, sex, and body mass index (BMI). Spearman correlations were also conducted between baseline indole abundance/ratio and participant demographic factors (age, BMI, height, and weight). Additionally, sex differences in baseline indole abundance/ratio were tested using Mann-Whitney *U* tests and fold changes in median abundance/ratio between groups.

To investigate the potential effects of treatment on indoles, changes in indole abundance from pre- to post-treatment were tested using Wilcoxon signed-rank tests and fold changes. Partial Spearman rank correlations were conducted between post-treatment indole abundance/ratio and post-treatment HAM-D 17-item total score, HAM-A total score, and HAM-A Psychic and Somatic subscores, QIDS-SR 16-item total score, and each individual QIDS-SR item, after accounting for age, sex, and baseline BMI. The same analyses were also conducted with change from pre- to post-treatment scores of all psychiatric variables, and also with fold changes from pre- to post-treatment for each indole. Additionally, differences in indole post-treatment abundance/ratio and fold change were investigated between each pair of treatment response outcome groups (31) (treatment failure; partial response; response; remission) by conducting Mann-Whitney *U* tests. This analysis was also repeated with baseline indole abundances/ratios to investigate whether baseline levels of indoles may be associated with treatment outcome. All reported *p*-values were adjusted for multiple comparisons using the Holm method.

Neuroimaging analyses were conducted using AFNI (35) and jamovi (www.jamovi.org). Of 122 participants who had an adequate quality of resting-state fMRI data (32), 80 had metabolomic measurements and clinical scores. Voxel-wise linear regression analyses were performed to examine the relationship between SCC-FC and IS or psychic anxiety scores (uncorrected *p* < 0.005 and > 250 voxels cluster size). A conjunction analysis identified overlapping areas between the SCC-FC of IS and the SCC-FC of psychic anxiety scores. Subsequently, mediation analyses were performed using the Medmod module (43) in jamovi. Three regions identified by the whole brain linear regression analysis between SCC-FC and IS were used for the mediation analysis. We explored each region, and combinations of the three regions, in the mediation models. For each model, the direct and indirect effects were estimated using bootstrapping with 5000 samples.

## RESULTS

Of the 344 patients randomized in PReDICT, 196 had metabolomic measures available for analysis at baseline and 127 were available at week 12. The demographic and clinical characteristics of the 196 participants are presented in Table 1.

**Table 1.**
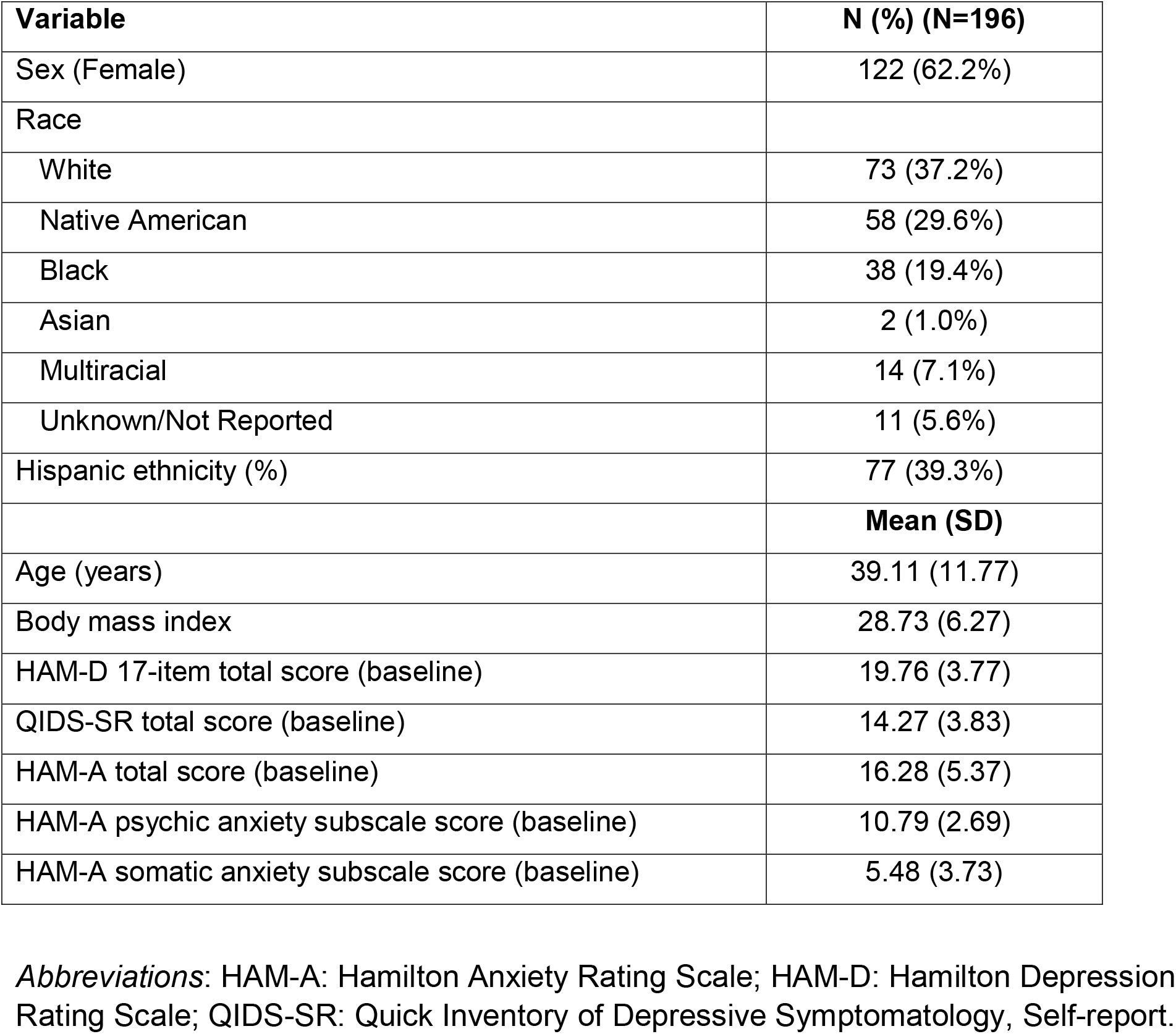
Subject demographic and clinical characteristics

### Baseline Associations

#### Associations of indole metabolites with demographic variables

Supplemental Figure 1 shows a heat map of correlations between baseline indole abundance/ratio and participant demographic variables. Abundance of ILA was positively associated with age, height, and weight (all *r*s > 0.18, all *p*s < 0.040). The ratios of IAA/ILA (negative associations) and IS/ILA (positive associations) were also significantly associated with height and weight (all *p*s < 0.034). For sex differences, abundance of IAA (Fold Changes (FC) = 1.19, *p* = 0.039) and ILA (FC = 1.37, *p* < 10^−9^), and ratios of ILA/IPA (FC = 1.40, *p* = 0.004) and ILA/IS (FC = 1.16, *p* = 0.013) were all found to be significantly higher in men than in women.

#### Associations of indole metabolites with depression and anxiety

Figure 2 shows a heat map of correlations between baseline indole abundance/ratio and baseline levels of the 17-item Hamilton Depression Rating Scale (HAM-D) (44) total score, Hamilton Anxiety Rating Scale (HAM-A) (45) total score, and HAM-A psychic and somatic subscores. Greater abundance of IS was associated with higher scores on the HAM-D 17-item total score (*r* = 0.21, *p* = 0.018), HAM-A total score (*r* = 0.26, *p* = 0.002), and HAM-A psychic subscore (*r* = 0.31, *p* = 0.0001), but not on the HAM-A somatic subscore. Additionally, the ratios of ILA/IS and IPA/IS were negatively correlated with HAM-A total and psychic scores (all *r*s > -0.20, all *p*s < 0.033), for which a negative correlation indicates that increasingly severe symptoms are associated with a relative increase in IS and/or a relative decrease in ILA or IPA. Additionally, IPA/IS was negatively correlated with HAM-D total score (*r* = -0.24, *p* = 0.001).

**Figure 2.**
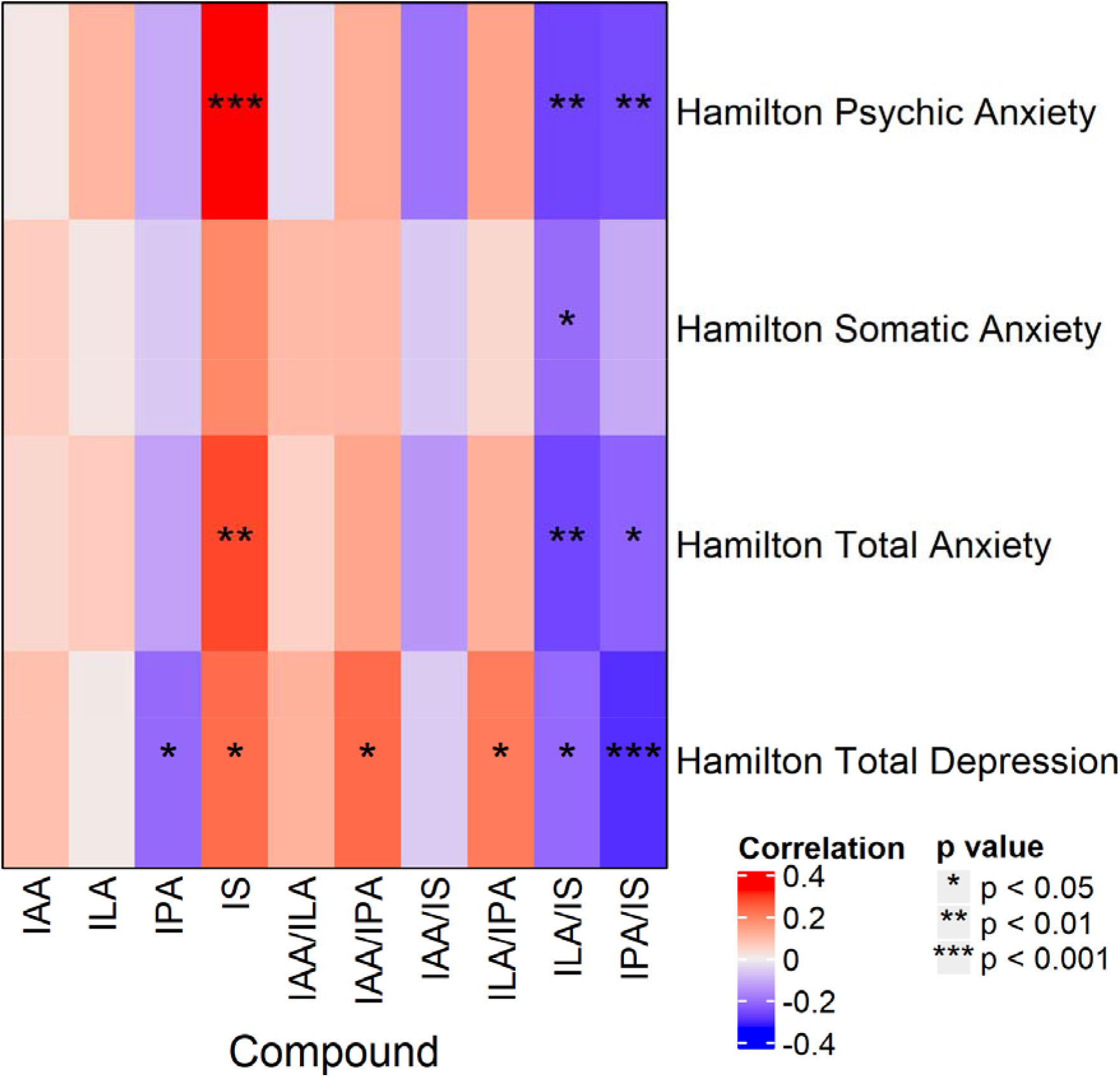
Heat map of partial Spearman rank correlations between baseline indole abundance/ratio and Hamilton Anxiety scores and Hamilton Depression scores, after accounting for age, sex, and BMI.

#### Associations of indole metabolites with individual symptoms of depression

Correlations between Quick Inventory of Depressive Symptoms – Self-Report (QIDS-SR) (34) items, and total scores and indole abundances/ratios are presented in Figure 3. Of note, IS positively correlated with items 4 (hypersomnia; *r* = 0.22, *p* = 0.016) and 6 (decreased appetite; *r* = 0.20, *p* = 0.034), and the IPA/IS ratio negatively correlated with QIDS-SR total score (*r* = -0.21, *p* = 0.027).

**Figure 3.**
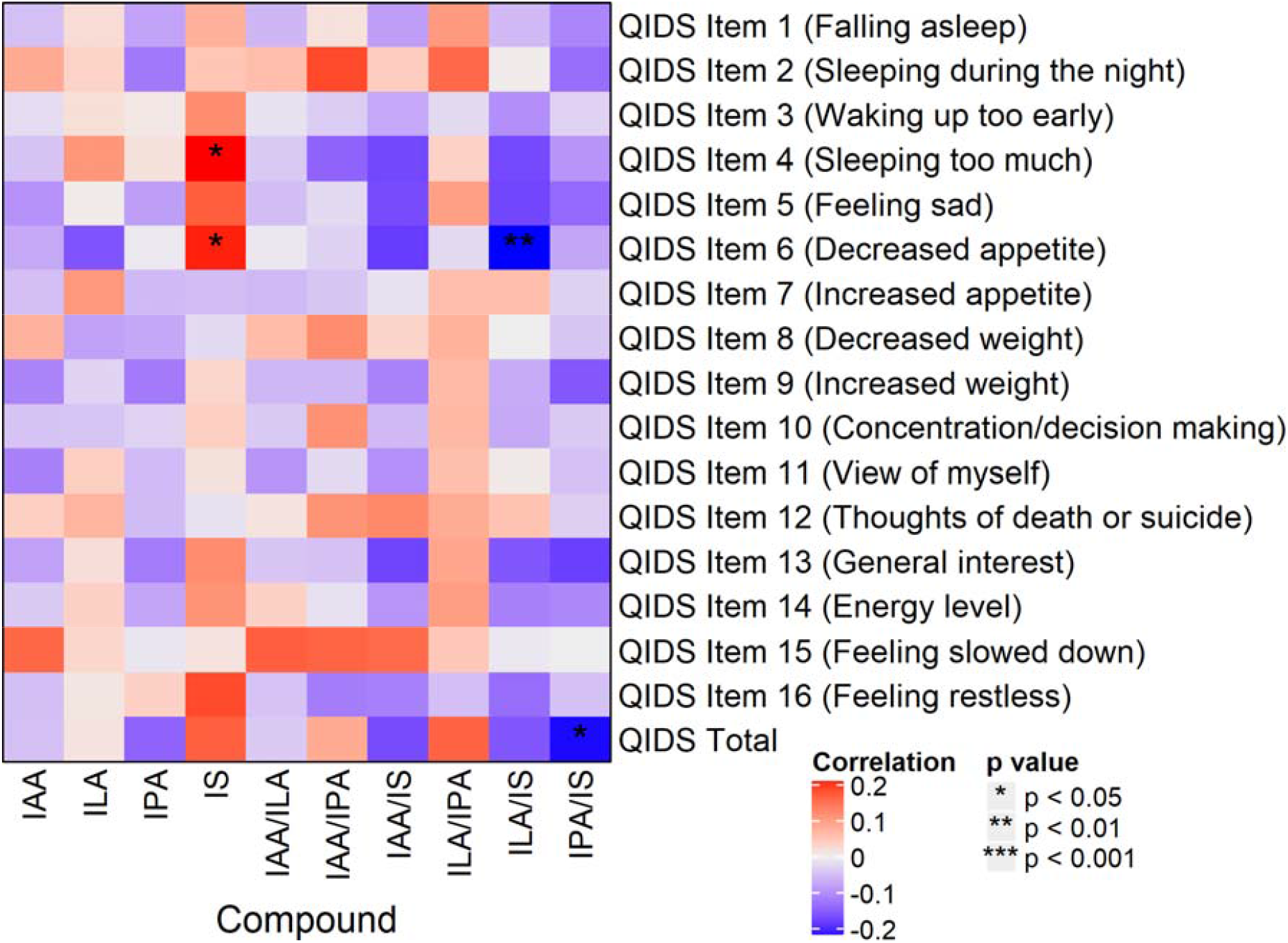
Heat map of partial Spearman rank correlations between baseline indole abundance/ratio and QIDS-SR items and total score, after accounting for age, sex, and BMI. *Abbreviations*: QIDS-SR: 16-item Quick Inventory of Depressive Symptomatology – Self-Rated.

### Treatment Effects

Compound abundance significantly increased from pre- to post-treatment for ILA (FC = 1.05, *p* = 0.006), but not for any other compound/ratio (all *p*s > 0.12). This indicates that the treatments had limited overall effect on indole composition and levels.

For post-treatment indole abundances and ratios, there were significant correlations between IAA/IS and QIDS-SR item 15 (feeling slowed down; *r* = 0.30, *p* = 0.007). Additionally, post-treatment ILA abundance was significantly higher in men than in women (FC = 1.24, *p* = 0.00001), as was the ILA/IPA ratio (FC = 1.54, *p* = 0.002). Conversely, the IPA/IS ratio was lower in men than in women (FC = 0.80, *p* = 0.048). No other significant post-treatment effects were observed.

For fold changes, change in IAA/IS ratio correlated with post-treatment scores of QIDS-SR item 5 (feeling sad; *r* = 0.27, *p* = 0.032). No significant associations were observed when correlating indole fold changes with any other post-treatment scores, or with change in depression/anxiety scores (all *p*s > 0.10). Additionally, no sex differences were observed for fold changes (all *p*s > 0.07), and no differences in fold changes were observed between response outcome groups (all *p*s > 0.12).

Baseline levels of indoles and their ratios did not significantly correlate with changes in symptoms for any measure or item (all *r*s < 0.15, *p*s > 0.59). Comparison of categorical response outcomes also showed no meaningful differences in baseline indole abundances or ratios. These analyses indicate that pre-treatment indole compound abundances are not predictive of eventual treatment outcomes.

#### Associations of indole metabolites with brain resting state functional connectivity

Relationships of Subcallosal Cingulate Cortex – Functional Connectivity (SCC-FC) with IS and with psychic anxiety scores are shown in Figure 4. IS abundance was positively correlated with SCC-FC with the bilateral anterior insula, anterior midcingulate cortex (aMCC), supplementary motor area (SMA), and right premotor area (Figure 4A). Psychic anxiety scores showed a significant positive correlation with SCC-FC with the left aMCC, right precuneus, and right premotor area; there was a negative correlation with SCC-FC with the ventromedial prefrontal cortex, right orbitofrontal, and left Brodmann Area 47 (Figure 4B). The conjunction analysis identified one overlapping area: the right premotor region (Figure 4C).

**Figure 4.**
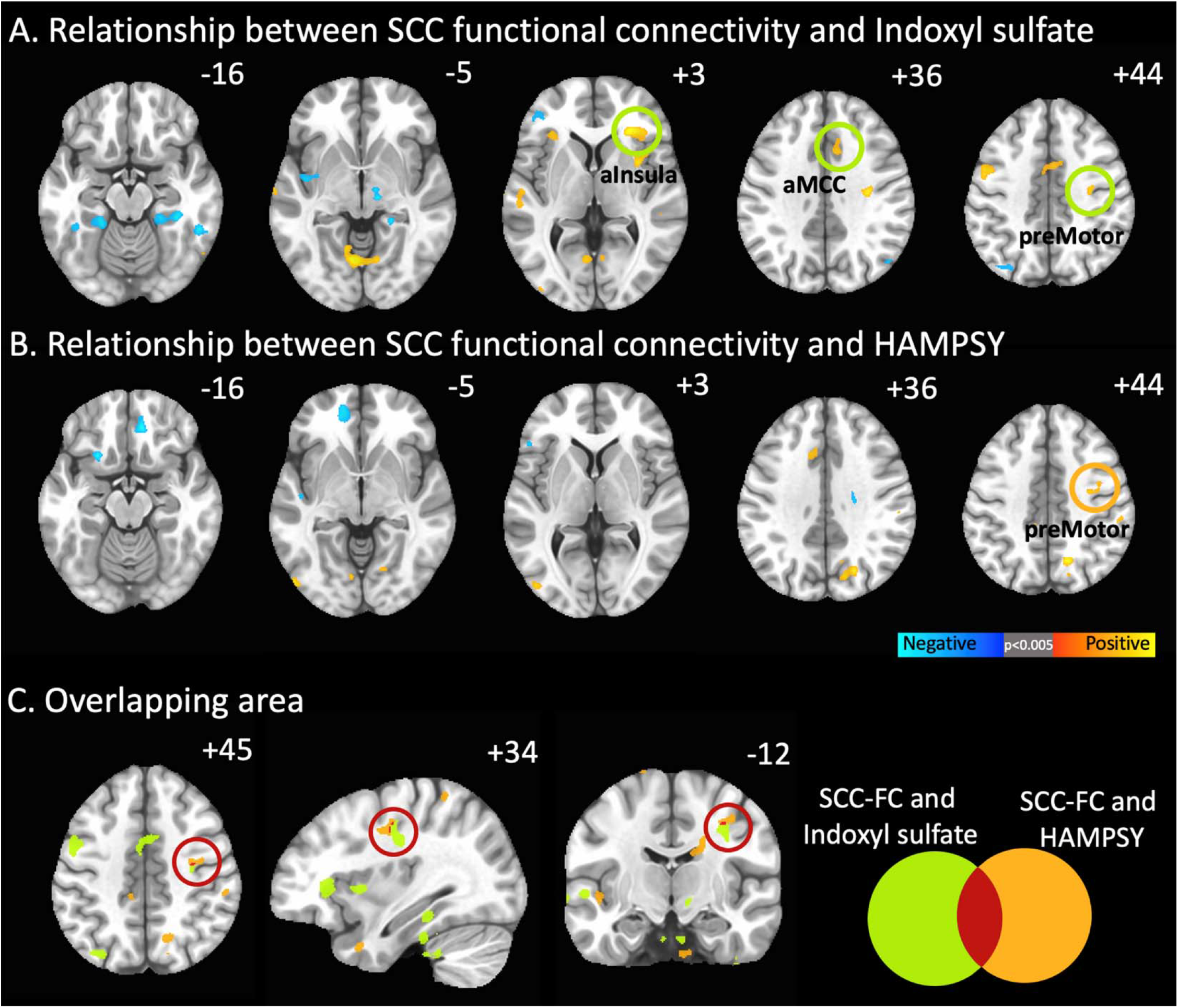
Resting state functional connectivity of subcallosal cingulate cortex (SCC) associations with peripheral indoxyl sulfate abundances and psychic anxiety scores. **A:** SCC functionally connected regions showing a significant correlation with indoxyl sulfate abundances. Orange circles identify regions incorporated into the mediation models. **B:** SCC functionally connected regions showing a significant correlation with psychic anxiety scores. Green circle identifies right premotor region. **C:** Conjunction analysis: SCC functionally connected region showing a significant correlation with both indoxyl sulfate abundances and psychic anxiety scores. The red circle indicates the only region to emerge in this analysis, the right premotor region. *Abbreviations*: HAMPSY: Psychic anxiety subscore of the Hamilton Anxiety Rating Scale. SCC-FC: subcallosal cingulate cortex functional connectivity

The mediation analyses explored whether the association of IS with psychic anxiety was mediated through its effects on SCC-FC. Figure 5A shows the overall association between IS and psychic anxiety (z = 1.976, *p* = 0.048). Figure 5B shows that the identified overlapping area in the SCC-FC analyses – the right premotor region – mediated the association between IS and psychic anxiety (indirect pathway: z = 2.138, *p* = 0.033). Because our whole brain SCC-FC analyses had also found IS concentrations to be significantly associated with two other regions previously identified in neuroimaging studies of anxiety (the right anterior insula and the aMCC, Figure 4A), we conducted further mediation analyses incorporating these two regions along with the right premotor region. Even though the three regions were highly correlated with each other in their functional connectivity to SCC (Figure 5C), only the right premotor region mediated the relationship between IS and psychic anxiety scores when all three regions were included in the model (Figure 5D, indirect pathway: z = 1.991, *p* = 0.046).

**Figure 5.**
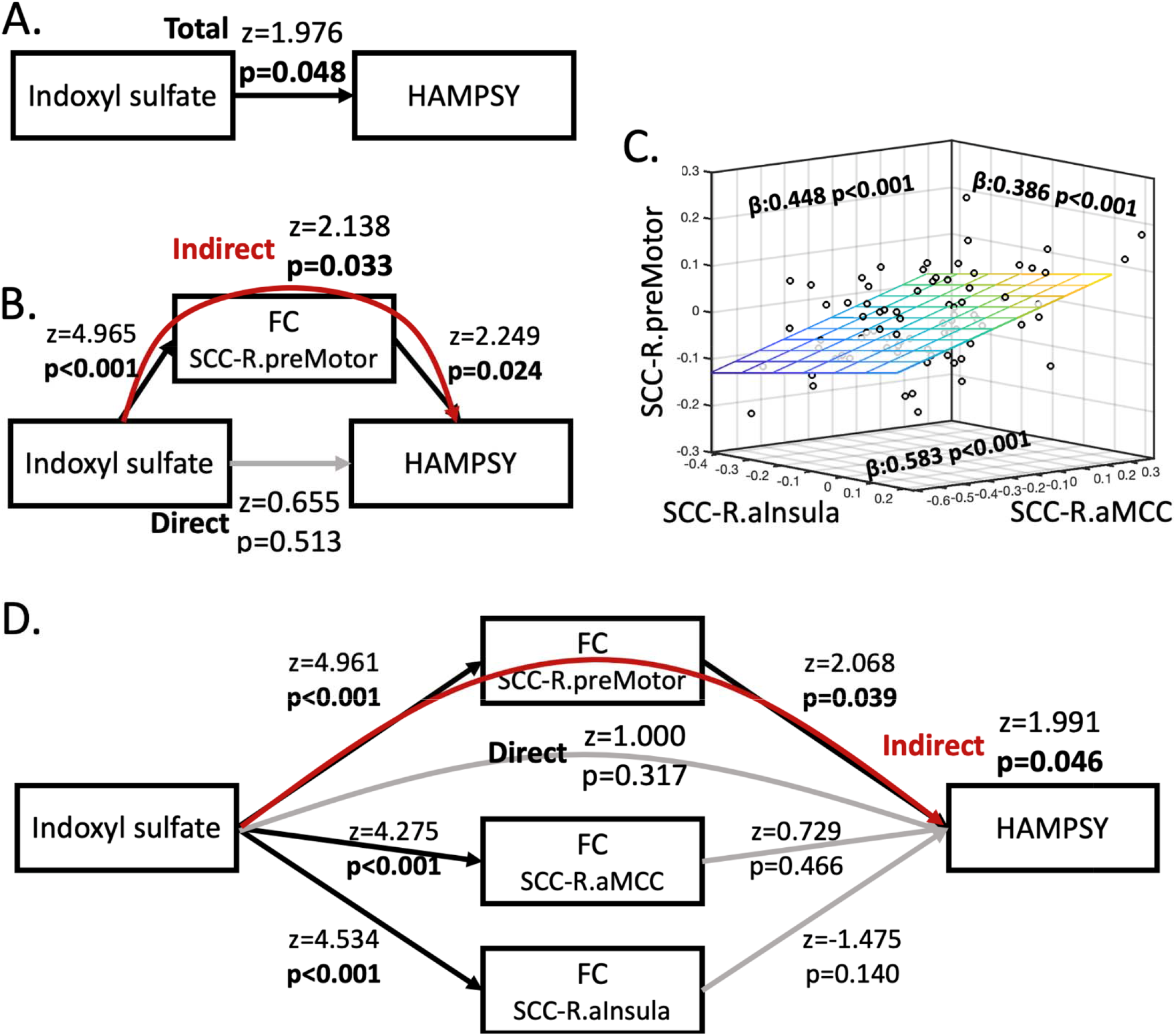
The impact of indoxyl sulfate on psychic anxiety scores is mediated by its effects on the resting state functional connectivity between the subcallosal cingulate cortex and the right premotor region. A. Association between indoxyl sulfate and psychic anxiety scores. B. Mediation model incorporating the overlapping area, right premotor region, indicating that the effect of indoxyl sulfate on psychic anxiety is mediated via its effects on the functional connectivity between the SCC and right premotor region. C. Significant SCC-FC correlations between the right anterior insula, right anterior midcingulate cortex, and right premotor region, which were included in the mediation model shown in D. D. Full mediation model incorporating the three regions showing significant SCC-FC correlations with indoxyl sulfate abundances. Although indoxyl sulfate is significantly correlated with all three regions, only the pathway through the right premotor region significantly mediates indoxyl sulfate’s effect on psychic anxiety. *Note*: Black lines indicate significant associations within the model; grey lines are insignificant associations. Red line indicates significant mediation of indoxyl sulfate on psychic anxiety through the indirect pathway of right premotor SCC-FC. *Abbreviations*: HAMPSY: Psychic anxiety subscore of the Hamilton Anxiety Rating Scale. SCC-FC: subcallosal cingulate cortex functional connectivity.

## DISCUSSION

Increasing evidence suggests that gut bacteria can complement human metabolism, and that together they define the metabolome comprised of the collection of small molecules in blood and in different organs. Bacteria can further metabolize compounds available through human metabolism, food-intake, and/or human ingestion of chemicals. Also, humans can further metabolize compounds produced by bacteria, which results in human-bacteria co-metabolism and the production of a large number of chemicals that can impact human health, including brain function. Examples include the metabolism of cholesterol and its clearance mediated by bacteria, which can produce secondary bile acids that we recently implicated in the pathogenesis of Alzheimer’s disease (46,47). Several compounds produced from the metabolism of phospholipids and choline by gut bacteria lead to compounds like trimethylamine N-oxide, which have been implicated in cardiovascular diabetes and CNS disease (48,49).

Indoles represent a class of gut bacterially-derived compounds that are produced from tryptophan, an essential amino acid that can also be converted (through separate pathways) into tryptamine, serotonin, skatol, and melatonin, among other metabolites involved in CNS functioning and diseases. Mounting evidence suggests that indoles derived from gut bacterial metabolism exert significant biological effects and may contribute to the etiology of cardiovascular, metabolic, and psychiatric diseases. To date, research in this area has been mainly limited to experimental studies in model systems.

In this investigation, we interrogated levels of four indoles produced by gut bacteria and their relationship to anxiety and depression severity and response to treatment. At baseline, IS abundance was found to positively correlate with severity of Psychic Anxiety and total anxiety. IPA seems protective, as noted earlier, indicating that indoles, as a class, can have mixed effects on neuropsychiatric health. Different strains of bacteria can lead to the production of different indoles; for example, tryptophanase-producing bacteria produce toxic IS while *Clostridium sporogenes* and other bacterial strains produce protective IPA (50,51). Notably, IS levels did not meaningfully change with treatment, and changes in IS were not correlated with improvement in depression or anxiety measures. This suggests that gut microbiome composition and activity might be modulated as an approach for developing additional classes of therapies effective in the treatment of anxiety and depression.

This association between IS and anxiety seems to be mediated by the impact of IS on the functional connectivity between the SCC and the right premotor region. IS abundances were also associated with activation of a well-established network for the processing and control of emotionally salient, particularly aversive, stimuli, comprising the anterior insula and aMCC. Taken together, these results suggest that the co-metabolism of tryptophan by certain gut microbiota that result in the production of IS, which can induce anxiety through the activation of established brain networks, and that existing treatments do not specifically resolve this pathogenetic process when they lead to clinical improvement.

The neuroimaging analyses indicate that the effect of IS on psychic anxiety symptoms is mediated through the functional connectivity of the SCC with the premotor cortex, as part of a network involved in processing emotionally salient stimuli. The anterior insula, aMCC, and supplementary motor area form a network that is involved in the attention to, interpretation of, and control of emotional responses (52-56). The premotor cortex is functionally and structurally connected to the SMA and the aMCC, which act together in the preparation and readiness for voluntary movement in response to internal and external stimuli (57), and the aMCC is a site of integration for the processing of pain and motor control (58). Outputs from this network include projections to the spinal cord and adrenal medulla (59), which may contribute to the sympathetic arousal and heightened cortisol release under situations of psychic stress.

Although the activity of the premotor cortex has not been a major focus in studies of anxiety and depression, Pierson and colleagues (60), using the electroencephalography measure of contingent negative variation (which localizes to the premotor cortex (61)), demonstrated abnormal activation of this region in anxious MDD patients compared to MDD patients with psychomotor retardation. Ma and colleagues (62) found that patients with generalized anxiety disorder have increased resting state functional connectivity between the habenula and right premotor cortex. Others have found abnormal premotor function in social anxiety disorder (63).

Our finding of an association between IS and activation of the insula bilaterally is consistent with the insula’s known involvement in processes relevant to anxiety, including emotional salience (64), empathy for others’ pain, and processing of uncertainty (55,65-67). In contrast, we did not find an association between IS abundances and somatic anxiety scores, nor was there an association of IS abundances with functional connectivity of the SCC-posterior insula, the insular region involved in sensorimotor integration. This reveals the specificity of the IS-anterior insula association for psychic anxiety.

Conceptualizing psychic anxiety as a chronic aversive stimulus akin to long-term pain may explain the positive correlation between higher IS levels and the QIDS-SR loss of appetite item. In mice, inflammatory pain is inhibited in the presence of hunger, mediated by neuropeptide Y signaling in the parabrachial nucleus (68). The association of higher IS concentrations with both reduction in appetite and increased connectivity between brain regions involved in pain processing (anterior insula and aMCC) may indicate that the symptom of low appetite reflects a compensatory response to this chronic anxiety-type pain.

Limitations of this study include the absence of a healthy control comparison group. We lacked fecal samples we could analyze which would allow for a more direct correlation between specific gut microbiome species and the IS measures. We could not determine whether IS is the etiological agent of the anxiety because IS also acts to reduce the integrity of the blood brain barrier (28), thereby creating the possibility that CNS penetration by an alternative molecule in the periphery is responsible for the observed association between anxiety and IS.

Taken together, our results indicate that increases in IS lead to the activation of an established network that is involved in the processing and control of aversive stimuli, but that the conscious experience of anxiety depends upon the degree of IS-related activation of SCC-right premotor cortex functional connectivity. The absence of an association between psychic anxiety scores and anterior insula/aMCC SCC-FC (Figure 4B) may indicate that although IS activates this control network in all patients, it is only when network function is inadequate that psychic anxiety ensues in conjunction with premotor activation in preparation for action (69). These analyses reveal the potential of integrated peripheral metabolomic-neuroimaging analyses to reveal mechanistic pathways that are associated with neuropsychiatric symptoms, especially for characterizing the pathological impact of specific gut microbiome-derived metabolites.

## Acknowledgements

We acknowledge the editorial services of Mr. Jon Kilner, MS, MA (Pittsburgh) and the assistance of Ms. Lisa Howerton (Duke). This work was funded by grant support to Dr. Rima Kaddurah-Daouk (PI) through NIH grants R01MH108348, R01AG046171 and U01AG061359. Dr. Boadie Dunlop has support from NIH grants P50-MH077083 (PI Mayberg), R01-MH080880 (PI Craighead), UL1-RR025008 (PI Stevens), M01-RR0039 (PI Stevens) and the Fuqua Family Foundations.

## Disclosures

Dr. Dunlop has received research support from Acadia, Compass, Aptinyx, NIMH, Sage, and Takeda, and has served as a consultant to Greenwich Biosciences, Myriad Neuroscience, Otsuka, Sage, and Sophren Therapeutics.

Dr. Rush has received consulting fees from Compass Inc., Curbstone Consultant LLC, Emmes Corp., Holmusk, Johnson and Johnson (Janssen), Liva-Nova, Neurocrine Biosciences Inc., Otsuka-US, Sunovion; speaking fees from Liva-Nova, Johnson and Johnson (Janssen); and royalties from Guilford Press and the University of Texas Southwestern Medical Center, Dallas, TX (for the Inventory of Depressive Symptoms and its derivatives). He is also named co-inventor on two patents: U.S. Patent No. 7,795,033: Methods to Predict the Outcome of Treatment with Antidepressant Medication, Inventors: McMahon FJ, Laje G, Manji H, Rush AJ, Paddock S, Wilson AS; and U.S. Patent No. 7,906,283: Methods to Identify Patients at Risk of Developing Adverse Events During Treatment with Antidepressant Medication, Inventors: McMahon FJ, Laje G, Manji H, Rush AJ, Paddock S. Dr. Kaddurah-Daouk in an inventor on a series of patents on use of metabolomics for the diagnosis and treatment of CNS diseases and holds equity in Metabolon Inc.

## Author Contributions

CRB did analysis of data and helped write the manuscript; OF and his team generated biochemical data and wrote its methods and helped with interpretation of findings; HSM, WEC, JC, KSC and BWD did analysis connecting metabolomics data to imaging data and helped with writing of manuscript; SMD, SB and HS helped with background literature searches and with interpretation of findings; RRK, BWD and AJR helped with interpretation of findings and clinical relevance; RKD is PI for project helped with concept development, study design, data interpretation and connecting biochemical and clinical data, and with writing of manuscript.

