## Supplemental Data for "Indoxyl sulfate, a gut microbiome-derived uremic toxin, is associated with psychic anxiety and its functional magnetic resonance imaging-based neurologic signature"

**Supplemental Figure 1.** Heat map of Spearman rank correlations between baseline indole intensity/ratio and participant demographic variables.

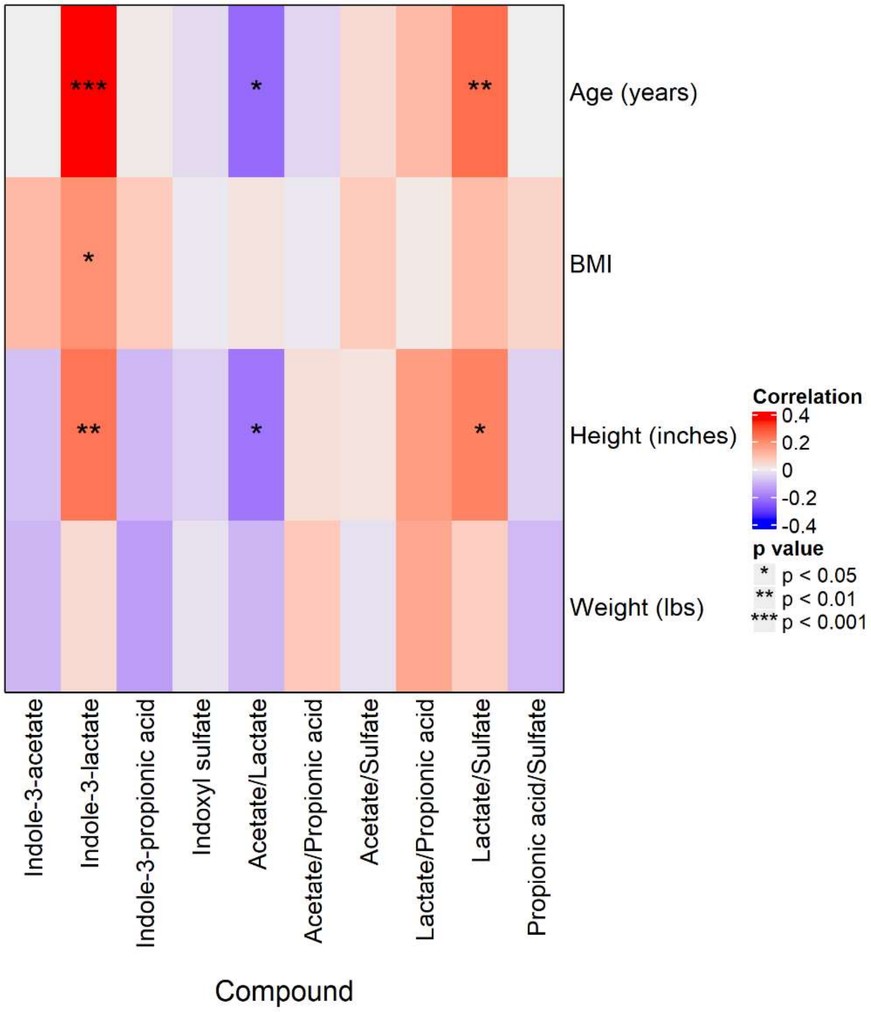
